# Potent Neutralizing Activity of Polyclonal Equine Antibodies against Omicron SARS-CoV-2

**DOI:** 10.1101/2022.02.21.481341

**Authors:** J. Luczkowiak, P. Radreau, L. Nguyen, N. Labiod, F. Lasala, C.H. Herbreteau, R. Delgado

## Abstract

Using a polyclonal approach of equine anti-SARS-CoV-2 F(ab’)2 antibodies we have achieved a high level of neutralizing potency against all SARS-CoV-2 variants tested. Neutralization titers were in the range of 10^5^-10^6^ IU/mL including Omicron: 111,403 UI/mL, which is 2-3 orders of magnitude what is normally achieved in response to SARS-CoV-2 infection and/or vaccination. The presence of high titers of a repertoire of antibodies targeting conserved epitopes in different regions of the spike protein could plausibly account for this remarkable breadth of neutralization. These results warrant the clinical investigation of anti-SARS-CoV-2 equine polyclonal F(ab’)2 antibodies as a novel therapeutic strategy against COVID-19

## Background

The rapid widespread of SARS-CoV-2 during the COVID-19 pandemics has caused 400 M of confirmed cases and over 5 M deaths by February 2022(1) along with significant socio-economic disruptions. This active transmission of SARS-CoV-2 has resulted in the emergence of several variants of concern (VoC) that apparently have been selected by a higher transmissibility and have challenged the public health control strategies to contain the pandemics(2). The recent appearance of the variant Omicron, that combines an augmented transmission capability along with an evasion from neutralization by convalescent or vaccinees sera, is a further hurdle for pandemics control(3). A number of potent monoclonal antibodies (mAbs) have received emergency authorization by EMA and FDA for COVID-19 treatment in selected patients(4). However, most of these mAbs have been rendered inefficacious by the highly mutated Omicron variant(5).

Considering the experience on the evolution of SARS-CoV-2 into a diversity of variants, a polyclonal approach in which many potential epitopes within the SARS-CoV-2 Spike protein are targeted could have great advantages in terms of breadth to neutralize current and potential future SARS-CoV-2 VoC. On the other hand, heterologous immunoglobulins have been used for more than a century in human therapy (especially for envenomation, rabies and tetanus)(6). Specific polyclonal immunoglobulins are indeed a well-known and effective therapeutic alternative that can be used quickly to respond against major health risks such as pandemics, emerging diseases and bioterrorism. FBR-002 is composed of purified polyclonal equine fragments F(ab’)2 directed against the SARS-CoV-2 spike protein. FBR-002 represents a safe and efficient therapy candidate to treat COVID-19 hospitalized patients by i) its polyclonality that allows to target multiple epitopes of the spike protein, limiting the risk of viral escape if new strains emerge and ii) its highly purifying process and lack of Fc portion minimizing the risk of Antibody-Dependent-Enhancement (ADE), immunogenicity and overall side-effects pattern (eg serum sickness syndrome) compared to whole immunoglobulins that may be used for passive immunotherapy. Therefore, F(ab’)2 polyclonal fragments show less safety concerns as compared to other related products such as plasma from recovered patients or polyclonal humanized anti-SARS-CoV2 antibodies. In this report we described the *in vitro* evaluation of FBR-002, a clinical grade product of polyclonal F(ab’)2 immunoglobulin fragments from Fab’entech against SARS-CoV-2 VoC.

## Materials & Methods

### Equine hyperimmune plasma

Three healthy French trotter horses were hyperimmunized with full SARS-CoV-2 spike protein. Horses had no detectable antibodies against SARS-CoV-2 before immunization and were strictly controlled for several viruses. Blood samples were collected regularly after immunization and plasma were prepared and stored at -20°C.

### Preparation of F(ab’)2 fragments

Pooled horse plasma was purified as previously described to obtain highly purified F(ab’)2 fragments(7, 8). The main purification steps consist of: i) anion-exchange chromatography steps, which eliminate proteins including albumin; ii) hydrolysis of whole immunoglobulins into F(ab’)2 fragments in order to eliminate Fc fragments, which are horse specific; and iii) pasteurization (heat treatment) at 60°C for 10 h to ensure viral safety of the product. The final bulk product (FBR-002) is filtered at a level of 0.2 µm and then stored at +5°C ± 3°C until use. The FBR-002 batch used in this study was purified (>99.5%) under GMP conditions and prepared at a final concentration of 18 g/L.

### Cell lines

Baby hamster kidney cells (BHK-21/WI-2, Kerafast # EH1011, Boston, MA, USA) and African Green Monkey Cell Line (VeroE6) were cultured in Dulbecco’s modified Eagle medium (DMEM) supplemented with 10% heat-inactivated fetal bovine serum (FBS), 25 μg/ml gentamycin and 2 mM L-glutamine.

### Production of SARS-CoV-2 pseudotyped VSV

VSV-G pseudotyped replication deficient rVSV-luc recombinant viruses were produced following previously published protocol(9, 10). The SARS-CoV-2 Spike mutant D614G was generated by site-directed mutagenesis using as an input DNA the expression vector encoding SARS-CoV-2 Spike_614D protein (kindly provided by J. Garcia-Arriaza, CNB-CSIC) by Q5 Site Directed Mutagenesis Kit (New England Biolabs, Barcelona, Spain) following the manufacturer’s instructions. SARS-CoV-2 variant Alpha (B.1.1.7, GISAID: EPI_ISL_608430), SARS-CoV-2 variant Beta (B.1.351, GISAID: EPI_ISL_712096), SARS-CoV-2 variant Gamma (P.1, GISAID: EPI_ISL_833140), SARS-CoV-2 variant Delta (B.1.617.2, GISAID: EPI_ISL_1970335), SARS-CoV-2 variant Omicron (B.1.1.529, GISAID: EPI_ISL_6640917), SARS-CoV-2 variant Mu (B.1.621, GISAID: EPI_ISL_2828054), SARS-CoV-2 variant Kappa (B.1.617.1, GISAID: EPI_ISL_1970331), SARS-CoV-2 variant Iota (B.1.526, GISAID: EPI_ISL_1172950), SARS-CoV-2 variant Epsilon (B.1.427, GISAID: EPI_ISL_1969270) and MERS (GenBank: AFS88936.1) were synthesized and cloned into pcDNA3.1 by GeneArt technology (Thermo Fisher Scientific GENEART GmbH, Regensburg, Germany). SARS-CoV-1 was purchased from SinoBiological, China (Cat. No. VG40150-G-N). EBOV strain Makona GP (GeneBank: KM2330102.1) was synthesized and cloned into pcDNA3.1 by GeneArt® technology. BHK-21 were transfected to express the SARS-CoV-2 S protein using Lipofectamine 3000 (Thermo Fisher Scientific, Madrid, Spain) and after 24h transfected cells were inoculated with a replication-deficient rVSV-luc pseudotype (MOI: 3-5) that contains firefly luciferase instead of the VSV-G open reading frame, rVSVΔG-luciferase (G*ΔG-luciferase; Kerafast). After 1 h incubation at 37ºC, the inoculum was removed, cells were washed intensively with PBS and then the medium added. Pseudotyped particles were harvested 20 h post-inoculation, clarified from cellular debris by centrifugation and stored at -80ºC. Infectious titers were estimated as tissue culture infectious dose per mL by limiting dilution of the SARS-CoV-2 rVSV-luc-containing supernatants on Vero E6 cells. Luciferase activity was determined by luciferase assay (Steady-Glo® Luciferase Assay System, Promega, Madrid, Spain) in a GloMax® Navigator Microplate Luminometer (Promega, Madrid, Spain).

### Neutralization assay

A SARS-CoV-2-pseudotyped rVSV-luc system was used to test neutralizing activity. FBR-002, polyclonal F(ab’)2 immunoglobulin was tested in 3-6 replicates at dilutions 1:100, 500, 2,500, 12,500, 62,500, 312,500, 1,562,500. In addition to test sample, the neutralization assay included dilution series of an external calibrant with assigned unitage of 813 International Units per ml (IU/ml) for neutralization activity (WHO SARS-CoV-2 Serology International Standard, 20/136, Frederick National Laboratory for Cancer Research). For neutralization experiments, viruses-containing transfection supernatants were normalized for infectivity to an MOI of 0.5 and incubated with the dilutions of FBR-002 at 37º C for 1 h in 96-well plates. After the incubation time, 20,000 Vero E6 cells were seeded onto the virus-plasma mixture and incubated in a 37ºC for 24h. Cells were then lysed and assayed for luciferase expression. Neutralizing titer 50 (NT50) and 95% confidence intervals (95% CI) were calculated using a nonlinear regression model fit with settings for log inhibitor versus normalized response curves, in GraphPad Prism v8. Neutralization potency were calibrated using WHO International Standard 20/136. Calibrated NT50 in IU/ml were calculated as the observed NT50 titers multiplied by the calibration factor (assigned as 0.679), which is estimated as 813 IU/ml divided by NT50 (tested as 1:1,198) of the WHO International Standard 20/136(11). Ebola virus (EBOV) and Vesicular Stomatitis virus (VSV) rVSV-luc pseudotypes were used in parallel as controls at the same conditions as controls.

## Results

Results of the neutralization assays are summarized in the table 1 and figure 1 A and B. FBR-002 achieved a very high level of neutralizing potency against all SARS-CoV-2 variants tested: Neutralization titers (IU/mL) were in the range of 10^5^-10^6^. Specifically neutralizing titer against the Omicron VoC was 111,403. Although much lower, we observed FBR-002 neutralization against SARS-CoV-1 with titer of 13,232 IU/mL. There was no measurable neutralizing activity detected against MERS, EBOV or VSV.

**Table 1.** Neutralizing levels (IU/mL) with 95%CI against SARS-CoV-2: reference D614G, Alpha, Beta, Gamma, Delta, Omicron, Mu, Kappa, Iota, Epsilon, SARS-CoV-1, MERS, and controls: EBOV, VSV for FBR-002. CI: 95% Confidence Interval Nd: non detectable.

|  | **FBR-002** |  |
| --- | --- | --- |
|  | **IU/mL** | **95%CI** |
| **SARS-CoV-2 D614G** | 613369 | 526772 - 713925 |
| **SARS-CoV-2 Alpha** | 282449 | 240920 - 331136 |
| **SARS-CoV-2 Beta** | 340272 | 298966 - 387251 |
| **SARS-CoV-2 Gamma** | 437598 | 384853 - 497572 |
| **SARS-CoV-2 Delta** | 289484 | 230013 - 364287 |
| **SARS-CoV-2 Omicron** | 111403 | 91633 - 135448 |
| **SARS-CoV-2 Mu** | 489430 | 430485 - 556532 |
| **SARS-CoV-2 Kappa** | 295035 | 194628 - 447290 |
| **SARS-CoV-2 Iota** | 276126 | 219214 - 347817 |
| **SARS-CoV-2 Epsilon** | 228937 | 180033 - 291101 |
| **SARS-CoV-1** | 13232 | 8477 - 20646 |
| **MERS** | nd |  |
| **EBOV** | nd |  |
| **VSV** | nd |  |

**Figure 1.**
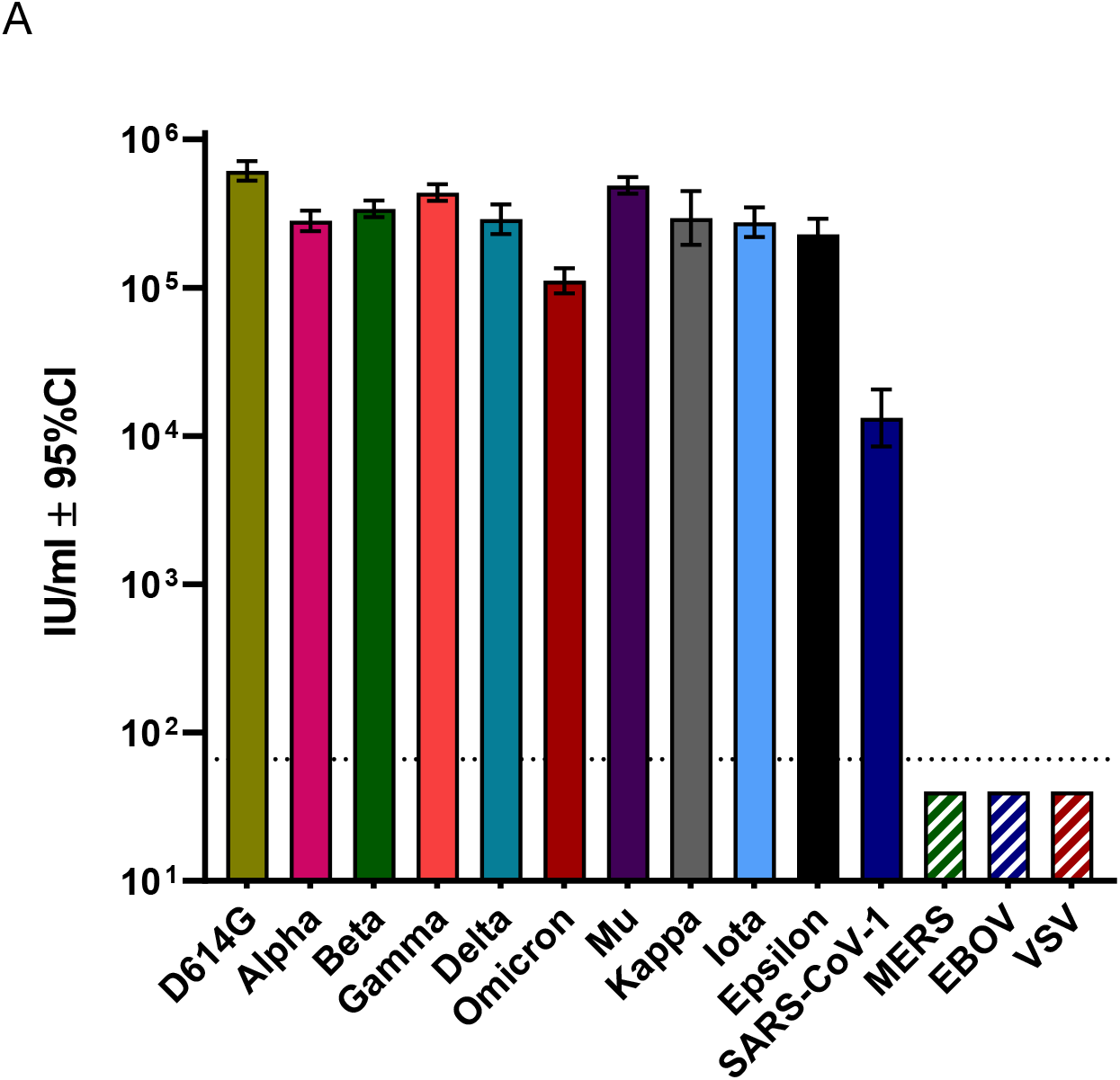
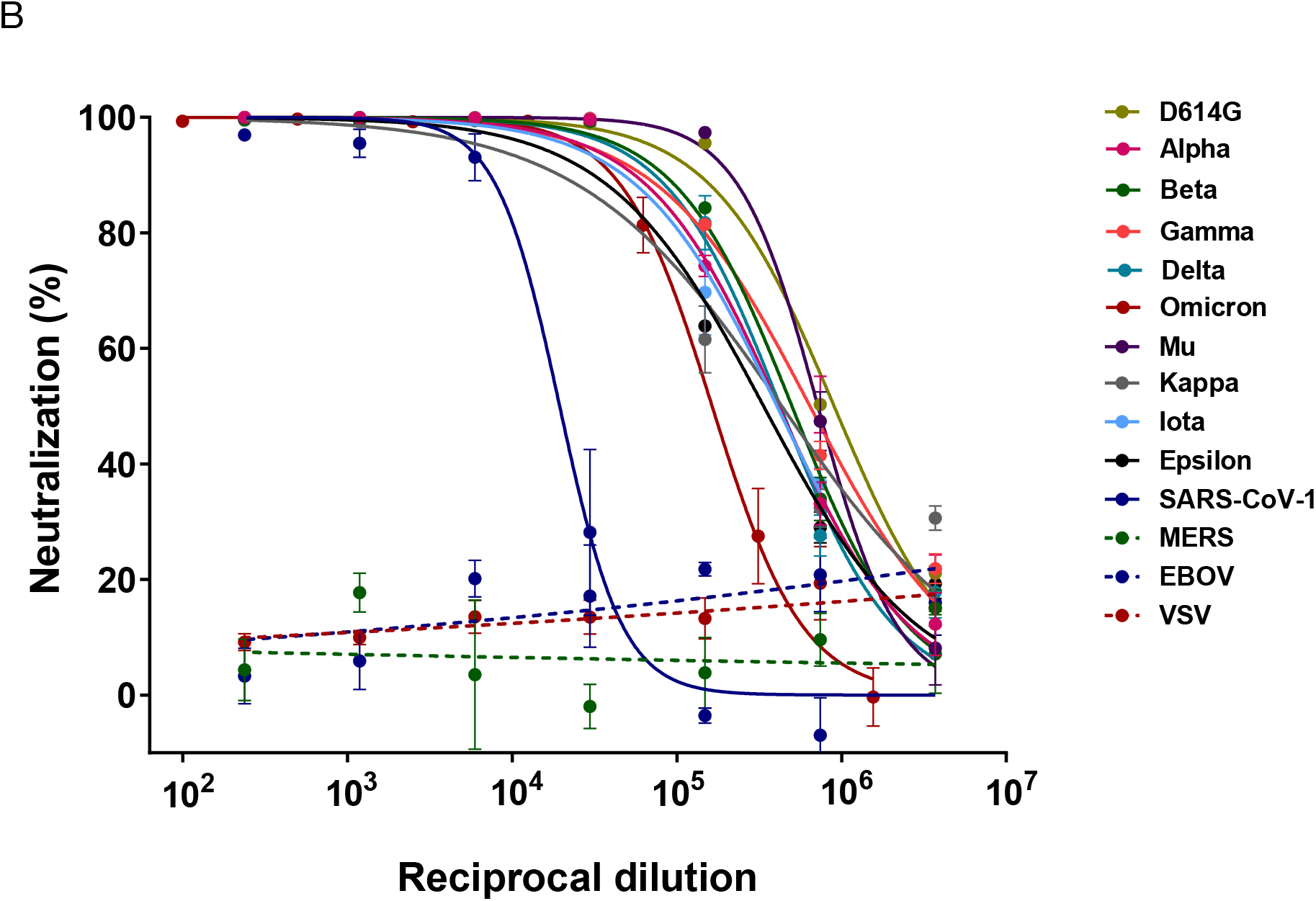
(A) SARS-CoV-2 neutralizing activity (NT50) of FBR-002 against SARS-CoV-2 VoC: reference D614G, Alpha, Beta, Gamma, Delta, Omicron, Mu, Kappa, Iota, Epsilon, SARS-CoV-1. Bars represent neutralizing IU/mL with 95% CI error bars. Calculated NT50 were calibrated using WHO International Standard 20/136. Dotted line neutralization positive threshold. (B) SARS-CoV-2 neutralizing activity curves for FBR-002 against SARS-CoV-2 VoC: reference D614G, Alpha, Beta, Gamma, Delta, Omicron, Mu, Kappa, Iota, Epsilon, SARS-CoV-1, MERS, EBOV and VSV. Neutralizing activity curves are presented using nonlinear regression model fit with settings for log inhibitor vs normalized response curve by GraphPad Prism V8.

## Discussion

The emergence of diversity along the evolution of the COVID-19 pandemics has led to the emergence of different SARS-CoV-2 variants whose adaptation properties to human transmission have resulted in an increased capability for transmission and in immune evasion to the neutralizing antibody response elicited by infection and or vaccination that is challenging the control of the pandemics. Among the antiviral strategies displayed, a number of monoclonal antibodies have been developed for therapeutic purposes. Monoclonal antibodies have been shown to be clinically effective for treatment of severe cases of emergent pathogens such as Ebola virus(12) and have been actively sought after for COVID-19. Most of the cloned neutralizing antibodies recognize epitopes within the Receptor Binding Motif (RBM) which is the contact region of the Receptor Binding Domain with the ACE2 receptor(13). These antibodies that recognize residues within the RBM are among the most common produced as response to SARS-CoV-2 infection in humans(14, 15). However, the evolution of scape mutations in the circulating VoC have rendered some of them inefficacious(16). This capability for immune evasion has been exhibited to a new limit by the recent emergence of the Omicron VoC that is resistant to neutralization by most of the clinically available monoclonal antibodies(17, 18).

Using a polyclonal approach of equine neutralizing anti-SARS-CoV-2 polyclonal F(ab’)2 antibodies we have achieved extraordinary neutralizing potency that is 2-3 orders of magnitude what is normally achieved in response to SARS-CoV-2 infection and/or vaccination(10). The neutralization coverage for the different variants is also remarkable, reaching >200000 IU/mL for most of VoC tested. As expected, Omicron showed the highest reduction in neutralization (5.5-fold as compared with the ancestral D614G sequence) however still was 111,403 IU/mL (Table 1, Figure 1 A and B). This neutralizing titer against Omicron is in fact two orders of magnitude higher that those obtained after booster vaccination in either naïve or COVID-19 convalescent individuals(19) and Luczkowiak J. et al (personal communication). Potency and breadth are properties of high titer polyclonal preparation that could be particularly helpful to target an evolving agent such as SARS-CoV-2, since considering the current level of transmission, even in highly immunized regions, the emergence of new variants can be expected. The presence of high titers of a repertoire of antibodies targeting conserved epitopes in different regions of the spike protein such as the receptor binding domain (RBD)(20) and also the N-terminal domain (NTD)(21) could plausible account for this remarkable breadth of neutralization. In this sense the neutralizing potency of the preparation is also significant against the related SARS-CoV-1 responsible for the SARS epidemic in 2002-3(22). These results warrant the clinical investigation of anti-SARS-CoV-2 equine polyclonal F(ab’)2 antibodies as a novel therapeutic strategy against COVID-19.

## Acknowledgments

This work was supported by the Instituto de Investigación Carlos III, ISCIII (grant numbers FIS PI1801007 and PI2100989); the European Commission Horizon 2020 Framework Programme (grant numbers 731868 project VIRUSCAN FETPROACT-2016, and 101046084 project EPIC-CROWN-2); and by Fundación Caixa-Health Research (grant number HR18-00469 project StopEbola).

## Notes

### Competing Interest Statement

Authors PR, LN and CHH are employees of Fab'entech, Lyon, France

### Summary of Updates

Names and affiliations

